# New Tricks for an old molecule: Preserved antibacterial activity of ribosomal protein S15 during evolution

**DOI:** 10.1101/2020.02.21.959346

**Authors:** Baozhen Qu, Zengyu Ma, Lan Yao, Zhan Gao, Shicui Zhang

**Affiliations:** Laboratory for Evolution & Development, Institute of Evolution & Marine Biodiversity and Department of Marine Biology, Ocean University of China, Qingdao 266003, China; Laboratory for Marine Biology and Biotechnology, Pilot National Laboratory for Marine Science and Technology (Qingdao), Qingdao 266003, China

## Abstract

Previous studies show that some ribosomal proteins possess antimicrobial peptide (AMP) activity. However, information as such remains rather fragmentary and limited. Here we demonstrated for the first time that amphioxus RPS15, BjRPS15, was a previously uncharacterized AMP, which was not only capable of identifying Gram-negative and -positive bacteria via interaction with LPS and LTA but also capable of killing the bacteria. We also showed that both the sequence and 3D structure of RPS15 and its prokaryotic homologs were highly conserved, suggesting its antibacterial activity is universal across widely separated taxa. Actually this was supported by the facts that the residues positioned at 45-67 formed the core region for the antimicrobial activity of BjRPS15, and its prokaryotic counterparts, including *Nitrospirae* RPS19_33-55_, *Aquificae* RPS19_33-55_ and *P. syringae* RPS19_50-72,_ similarly displayed antibacterial activities. BjRPS15 functioned by both interaction with bacterial membrane via LPS and LTA and membrane depolarization as well as induction of intracellular ROS. Moreover, we showed that RPS15 existed extracellularly in amphioxus, shrimp, zebrafish and mice, hinting it may play a critical role in systematic immunity in different animals. In addition, we found that neither BjRPS15 nor its truncated form BjRPS15_45-67_ were toxic to mammalian cells, making them promising lead molecules for the design of novel peptide antibiotics against bacteria. Collectively, these indicate that RPS15 is a new member of AMP with ancient origin and high conservation throughout evolution.

**Author summary:** Ribosomal protein, a component of ribonucleoprotein particles, is traditionally known involved in protein synthesis in a cell. Here we demonstrated for the first time that amphioxus ribosomal protein 15 was a novel antibacterial protein, capable of recognizing Gram-negative and -positive bacteria as well as killing them. It killed the bacteria by a combined mode of action of disrupting bacterial membrane integrity and inducing radical oxygen species production. We also showed that both eukaryotic ribosomal protein 15 and its prokaryotic counterpart ribosomal protein 19 possessed antibacterial activity, indicating that the antibacterial property is universal for this family of molecules. Moreover, we found that ribosomal protein 15 was present in the circulation system of various animals including shrimp, amphioxus, zebrafish and mice, suggesting it may physiologically play a key role in systematic immunity. Altogether, our study provides a new angle for understanding the biological function of ribosomal proteins.

## Introduction

The ribosome is an organelle within the cytoplasm of living cells that is composed of proteins and ribosomal RNAs (rRNAs), serving as the site for assembly of polypeptides encoded by messenger RNAs (mRNAs). Ribosomes are found in both prokaryotic and eukaryotic cells. In both types of cells, ribosomes are composed of two subunits, one large and one small [1, 2]. Each subunit has its own mix of proteins and rRNAs. The small and large subunits of eukaryotes are called 40S and 60S, respectively, while those of prokaryotes called 30S and 50S, separately. Ribosomal protein S15 (RPS15) is a component of the 40S subunit of eukaryotes, while its homolog in prokaryotes is S19 (RPS19) of the 30S subunit [3].

Ribosomal proteins, in addition to their conventional role in ribosome assembly and protein translation, are shown involved in diverse physiological and pathological processes, such as neurodegeneration in Parkinson’s disease, tumorigenesis, immune signaling and development [1,4,5]. Intriguingly, ribosomal proteins also show antimicrobial activity. Initially, the antimicrobial peptide (AMP) cecropin, first isolated from the moth *Hyalophora cecropia* [6–8], was mapped to the N-terminal region of the 50S ribosomal protein L1 of the pathogen *Helicobacter pylori* [9, 10]. Recently, the 50S ribosomal proteins L27 and L30 of the lactic acid bacterium *Lactobacillus salivarius* were shown to possess antimicrobial activity against *Streptococcus pyogenes*, *Streptococcus uberis* and *Enterococcus faecium* [11]. Furthermore, antibacterial activity was also observed for the 60S ribosomal protein L29 isolated from the gill of pacific oyster *Crassostrea gigas* [12] and the 40S ribosomal protein S30 isolated from the skin of rainbow trout *Oncorhynchus mykiss* [13]. It is thus clear that some ribosomal proteins of the small and large subunits of both prokaryotic and eukaryotic ribosomes can function as AMP. However, our information regarding ribosomal protein AMPs is rather fragmentary and limited. Moreover, little is known about the mode of action of ribosomal protein AMPs. In this study, we identified RPS15 of amphioxus (*Branchiostoma japonicum*), BjRPS15, as a novel member of AMP, and demonstrated that BjRPS15 executed its antimicrobial activity by both the interaction with bacterial membrane via LPS and LTA and membrane depolarization as well as production of intracellular ROS. We also showed that the emergence of antimicrobial activity of RPS15 could be traced to its prokaryotic homolog RPS19. This is the first report showing that RPS15 and its prokaryotic homolog RPS19 function as an AMP, much enriching our understanding of the biological activities of ribosomal proteins.

## Results

### Sequence characteristics and genomic structure of BjRPS15

The open reading frame (ORF) of *BjRPS15* (GenBank accession number: XP_019635827) obtained was 444 bp long, which encoded a deduced protein of 147 amino acids with a calculated molecular weight of about 16.96 kDa and an isoelectric point (pI) of 10.31 (Fig 1A). Analysis by SignalP showed that the deduced protein had no signal peptide, and analysis by SMART program revealed that the protein possessed a single Ribosomal-S19 domain at the residues 45 to 130 (Fig 1A). Analysis by Antimicrobial Peptide Calculator and Predictor at APD revealed that the BjRPS15 had a total hydrophobic ratio of 36% and a net charge of +19, suggesting that BjRPS15 is a putative AMP. Prediction by CAMP showed that the amino acid residues 45-67 (RRFSRGLKRKHLALIKKLRKAKK, designated BjRPS15_45-67_) were the core region of antimicrobial activity of BjRPS15 (Fig 1A). As shown in Table 1, the peptide BjRPS15_45-67_ had a total hydrophobic ratio of 34% and a net charge of +12.04. The 3D modeling revealed that BjRPS15 was composed of 7 α-helice and 3 β-sheets (Fig 1B), and BjRPS15_45-67_ comprised 2 α-helice (Fig 1C).

**Fig 1.**
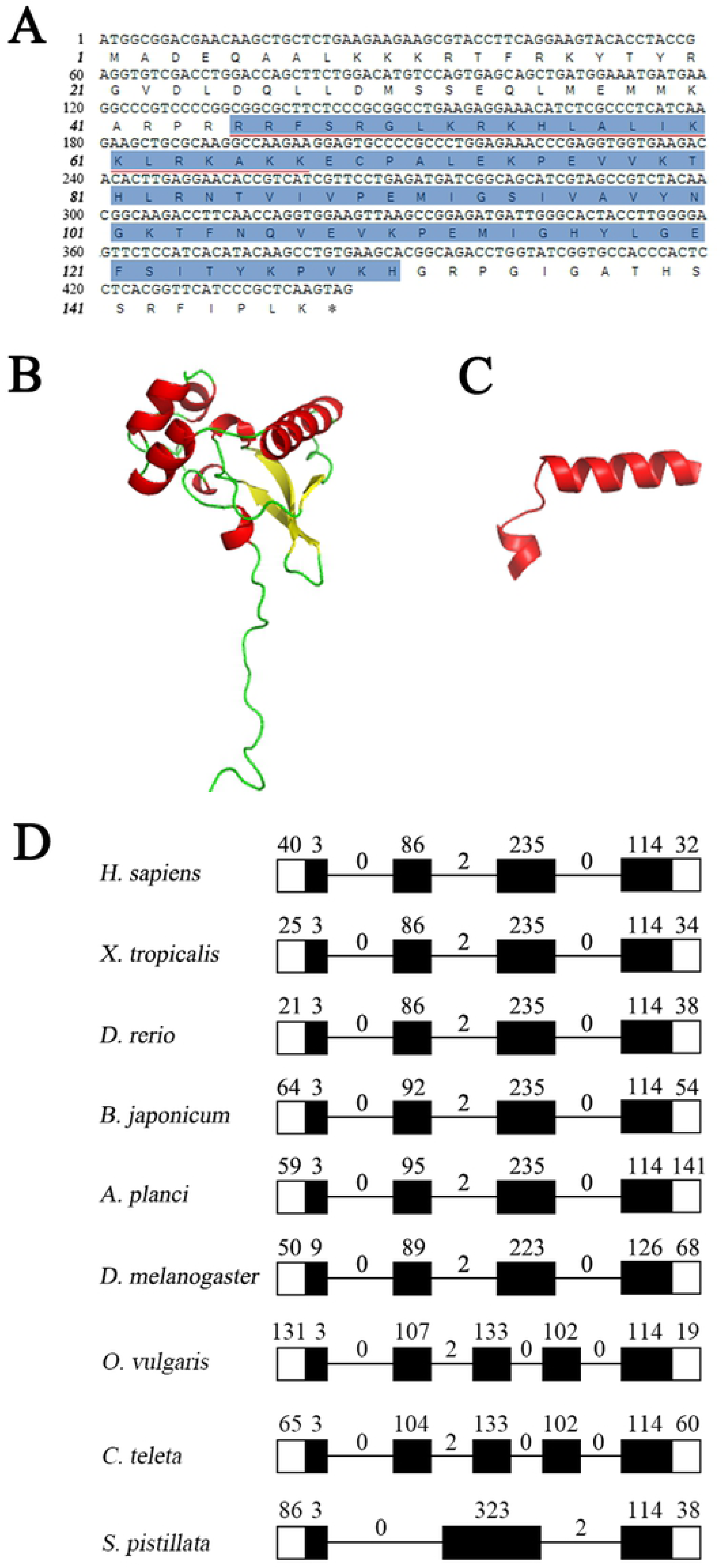
The full-length nucleotide and deduced amino acid sequences, 3D molecular modeling and genomic organization of BjRPS15. (A) The nucleotides and amino acids are numbered on the left margin. The termination codon is indicated with asterisk (*). The Ribosomal-S19 domain is shaded in blue. The peptide BjRPS15_45-67_ is underlined in red. (B) 3D structure model of rBjRPS15. (C) 3D structure model of BjRPS15_45-67_. (D) Exon-intron organizations of *BjRPS15* or *RPS15* in human, xenopus, zebrafish, starfish, fruit fly, octopus, Sea worm and coral. Exons are indicated with boxes and introns with lines; blacked boxes correspond to the coding regions and empty boxes to 5’ and 3’ untranslated regions. The length of exons and the phases of introns are shown. The accession numbers of gene used were listed in S2 Table.

**Table 1.**
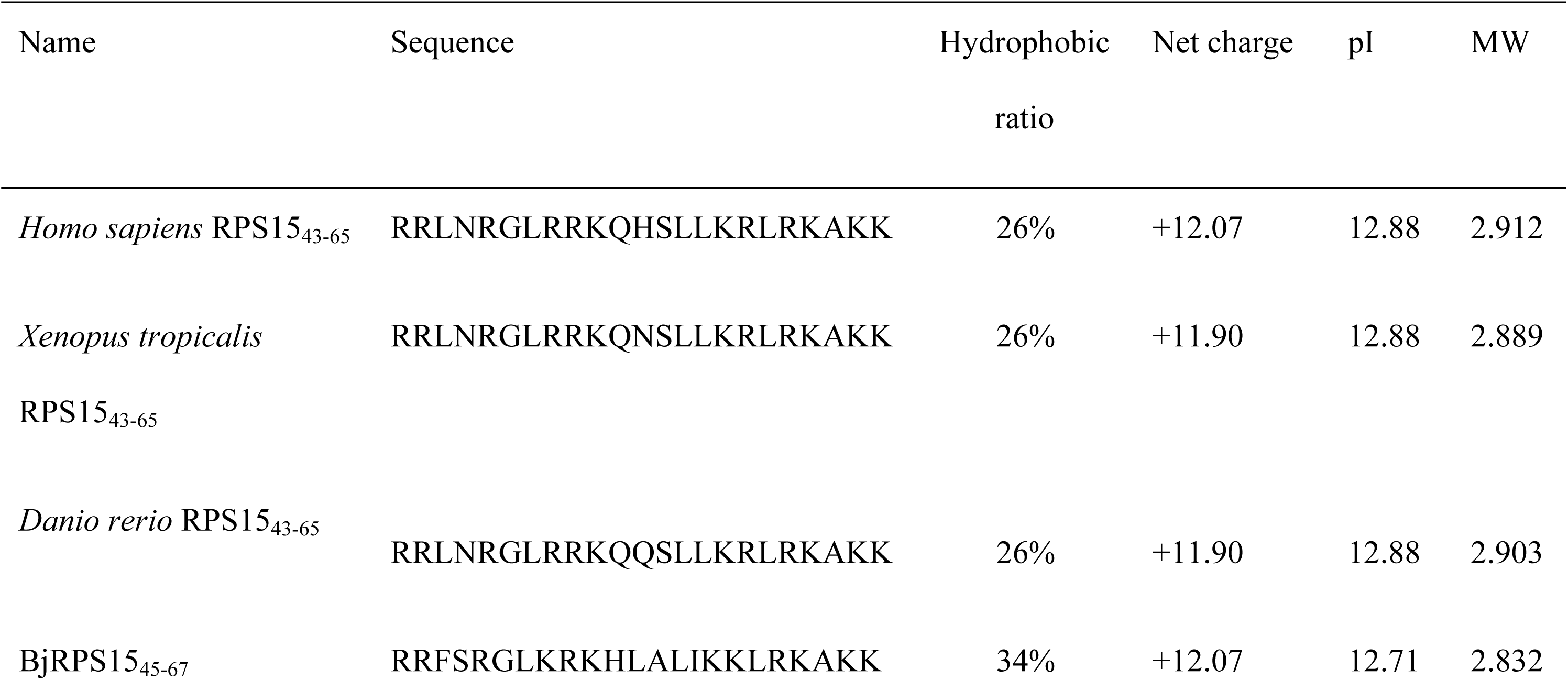
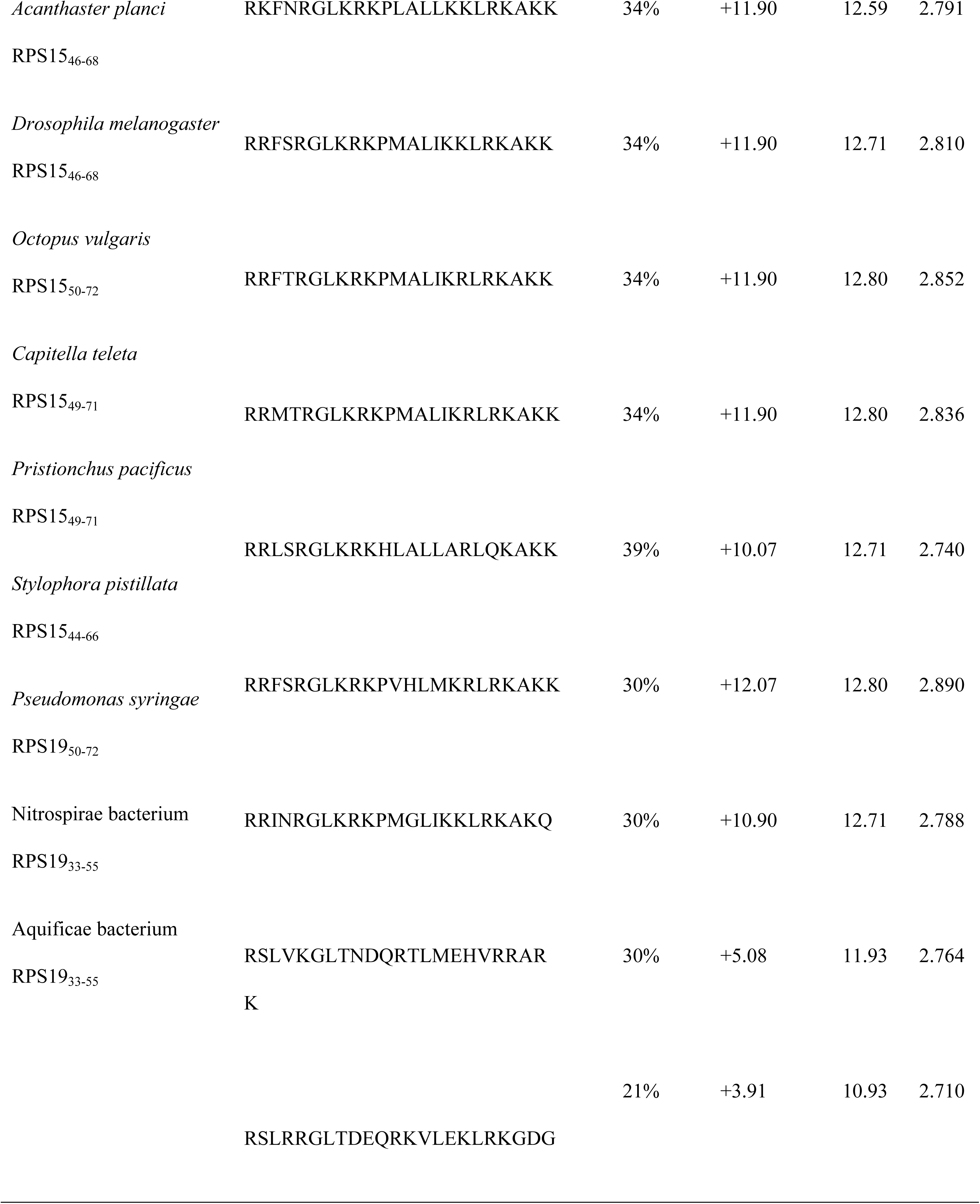
Amino acid sequences and chemical properties of BjRPS15 related peptides. pI, isoelectric point. MW, molecular weight (kDa).

Protein sequence comparison showed that BjRPS15 shared 75.2% to 78.6% identity to the RPS15 of vertebrates, including mammals, reptiles, amphibians and fishes [3,31,32], and 79.6% to 81.6% identity to that of invertebrates [3] as well as 41.2% to 73.9% identity to prokaryotic homolog RPS19 [33] (S1 Fig). A search of the completed draft assembly and automated annotation of amphioxus genomes revealed the presence of a single cDNA and its genomic DNA sequence in both *B. belcheri* and *B. floridae* (transcript id: XM_019780268.1 for *B. belcheri* and XM_002594975.1 for *B. floridae*). Both the cDNAs of *B. belcheri* and *B. floridae* encoded a protein with 100% identity to BjRPS15, suggesting that in amphioxus RPS15 was absolutely conserved in interspecies. Analysis of the genomic structure uncovered that all the homologs of *BjRPS15* from different animals comprised 3 to 5 exons interspaced by 2 to 4 introns (Fig 1D), but each of their coding exons shared highly identical sequence, suggesting the general genomic sequence of *RPS15* retained rather stable throughout multicellular animal evolution.

### Expression of *BjRPS15* after challenge with bacteria, LPS and LTA

qRT-PCR was used to examine the transcriptional profile of *BjRPS15* in the different tissues. As shown in Fig 2A, *BjRPS15* was predominantly expressed in the hepatic caecum, hind-gut, testis and ovary, and at a lower level in the gill, notochord and muscle, indicating that *BjRPS15* was expressed in a tissue-specific manner. Notably, the challenge with the Gram-negative bacteria *A. hydrophila* and *E. coli* and the Gram-positive bacteria *S. aureus* and *B. subtilis* both resulted in significant increase in the expression of *BjRPS15* (Fig 2B). Similarly, the challenge with LPS and LTA also induced marked increase in the expression of *BjRPS15* (Fig 2C). These data suggested that BjRPS15 might be involved in the anti-infectious response in amphioxus.

**Fig 2.**
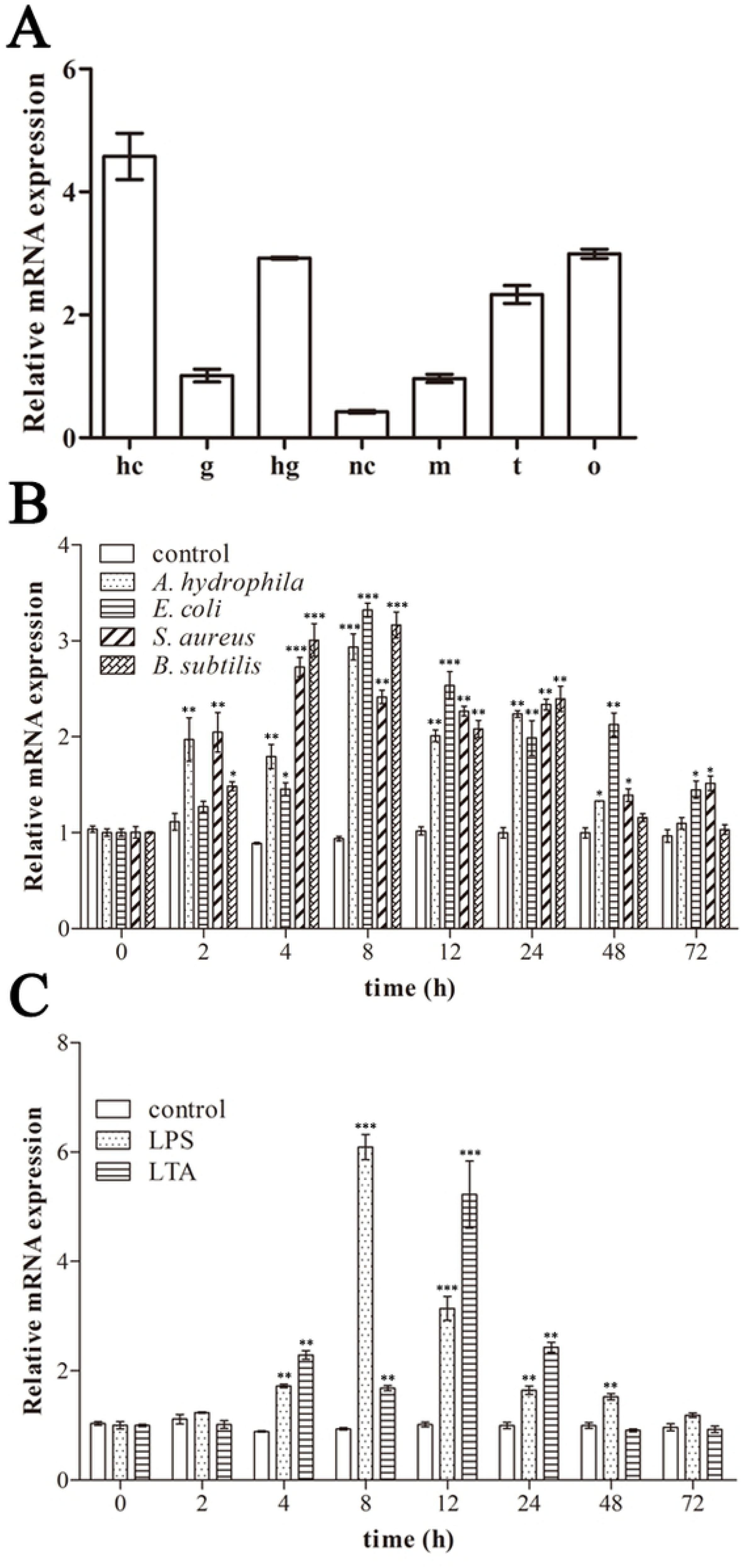
Transcriptional profiles of *BjRPS15* in different tissues (A), and in response to challenge with *A. hydrophila*, *E. coli*, *S. aureus* and *B. subtilis* (B) as well as LPS and LTA (C). The *EF1α* gene was chosen as internal control for normalization. (A) hc, hepatic caecum; g, gill; hg, hind-gut; nc, notochord; m, muscle; t, testis; o, ovary. (B and C) The animals were sampled at 0, 2, 4, 8, 12, 24, 48 and 72 h after the challenge, and total RNAs were extracted from the whole animals. The results shown are mean ± SEM, n = three replicates per group, and are pooled from three experiments per time point. Statistical differences between time points were assessed using unpaired Student’s t-test, \**p* < 0.05, \*\**p* < 0.01.

### Antibacterial activity of rBjRPS15 and BjRPS15_45-67_

The purified recombinant proteins rBjRPS15 and rTRX both yielded a single band of approximately 21.85 and 20.4 kDa, respectively, well matching the expected sizes (Fig 3A). Western blotting showed that rBjRPS15 and rTRX were both reactive with the anti-His-tag antibody (Fig 3A), indicating that they were properly expressed. We then tested the antimicrobial activity of rBjRPS15 and rTRX (control) against the Gram-negative bacteria *A. hydrophila* and *E. coli* as well as the Gram-positive bacteria *S. aureus* and *B. subtilis*. As shown in Fig 3B, rBjRPS15 showed conspicuous antimicrobial activities against all the bacteria tested, with the minimum bactericidal concentration MBC (defined as the lowest concentration at which the bacterium was completely killed) against *A. hydrophila*, *E. coli* and *B. subtilis* being about 2 μM and that against *S. aureus* being > 2 μM. We also evaluated the MBC_50_ (defined as the lowest concentration at which the 50% bacterium was killed) of rBjRPS15 against *A. hydrophila*, *E. coli, S. aureus* and *B. subtilis*, which was all about 0.5 μM. Similarly, BjRPS15_45-67_ also showed bactericidal activities against *A. hydrophila*, *E. coli*, *S. aureus* and *B. subtilis*, with the MBC_50_ against *A. hydrophila* and *E. coli* being about 2 μM and that against *S. aureus and B. subtilis* about 4 μM (Fig 3C). By contrast, rTRX showed little antimicrobial activity against all the bacteria tested (data not shown). These indicated that BjRPS15 was indeed an AMP with the residues 45-67 being the core region for the antimicrobial activity.

**Fig 3.**
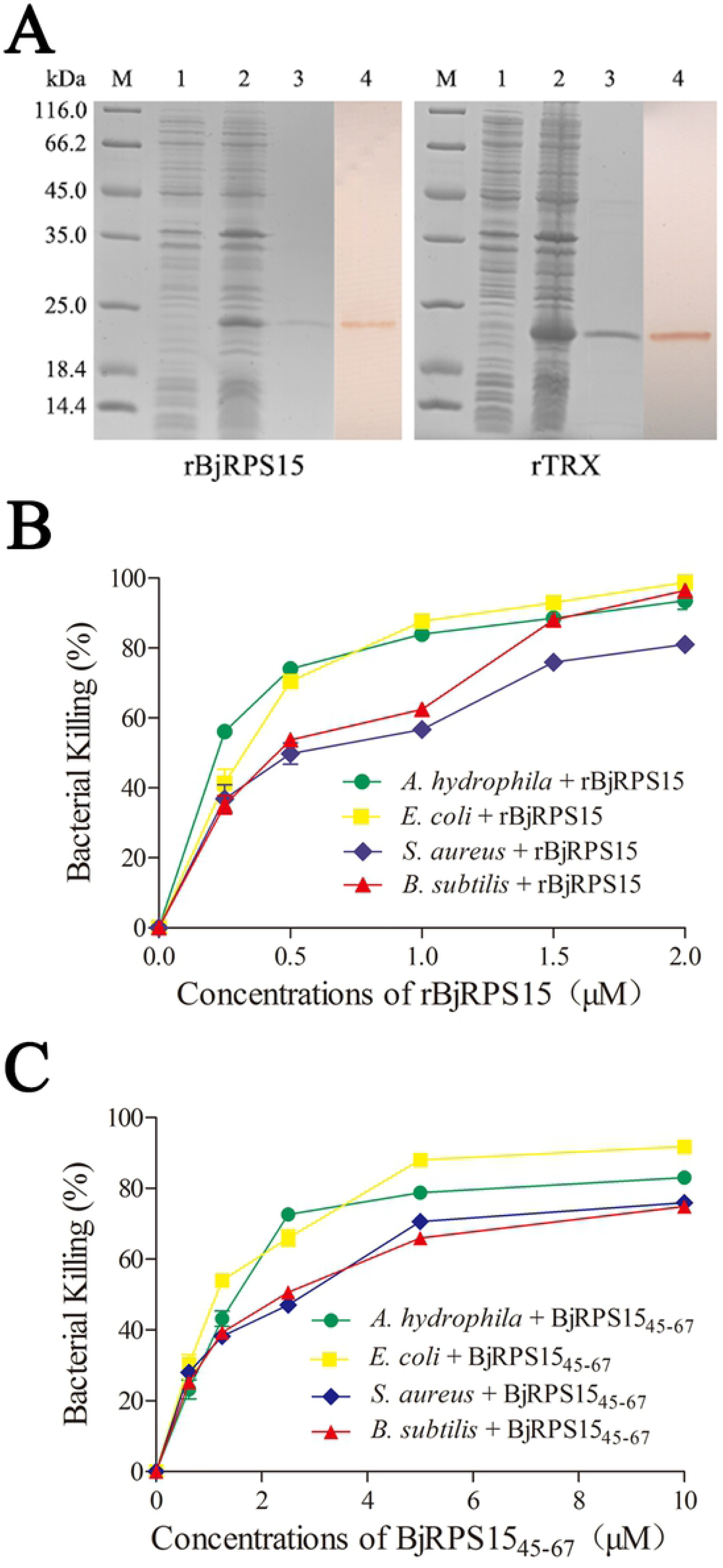
SDS-PAGE and Western blotting of recombinant proteins and antibacterial activity of recombinant rBjRPS15 and peptide BjRPS15_45-67_. (A) SDS-PAGE and Western blotting of recombinant proteins rBjRPS15 and rTRX. Lane M, marker; lane 1, total cellular extracts from *E. coli* transetta (DE3) containing expression vector before induction; lane 2, total cellular extracts from IPTG-induced *E. coli* transetta (DE3) containing expression vector; lane 3, purified recombinant proteins; lane 4, Western blot of purified recombinant proteins. The proteins on the gels were electroblotted onto PVDF membrane. After incubation with mouse anti-His-tag antibody, the membranes were incubated with the HRP-labeled goat anti-mouse IgG Ab and the bands were visualized using DAB kit. The concentrations of purified recombinant rBjRPS15 and rTRX were 100 μg/ml and 180 μg/ml respectively, and the amount of recombinant proteins loaded on the 12% SDS-PAGE gel was 20 μl. (B) Antibacterial activity of rBjRPS15 (0, 0.25, 0.5, 1 and 2 μM) and BjRPS15_45-67_ (0, 0.625, 1.25, 2.5, 5 and 10 μM) against bacteria. The rTRX and PBS were used as control.

### Antibacterial activity of BjRPS15_45-67_ counterparts

To test if the antimicrobial activity of RPS15 was conserved during evolution, the counterparts of BjRPS15_45-67_ ranging from prokaryotes to eukaryotes were investigated for the presence of antibacterial activity. First, sequence alignment showed that the sequence of BjRPS15_45-67_ was highly conserved among the eukaryotes as well as the prokaryotes, with the residues of RPS19_33-55_ of *Nitrospirae* sp. and RPS19_33-55_ of *Aquificae* sp. being most divergent (Fig 4). Analysis by Antimicrobial Peptide Calculator and Predictor at APD revealed that *Nitrospirae* RPS19_33-55_ had a hydrophobic ratio of 30% and a net charge of +5.08, *Aquificae* RPS19_33-55_ a hydrophobic ratio of 21% and a net charge of +3.91, and all the other BjRPS15_45-67_ counterparts a hydrophobic ratio of >25% and a net charge of +10 to +12.5 (Table 1). The 3D modeling showed that all the 3D structures of BjRPS15_45-67_ counterparts, including *Nitrospirae* RPS19_33-55_ and *Aquificae* RPS19_33-55_, were similar to that of BjRPS15_45-67_, consisting of 2 α-helice (S2 Fig). These suggested that the counterparts above might also have antibacterial activity. We thus synthesized the peptides of BjRPS15_45-67_ counterparts, including *H. sapiens* RPS15_43-65_, *X. tropicalis* RPS15_43-65_, *D. rerio* RPS15_43-65_, *A. planci* RPS15_46-68_, *D. melanogaster* RPS15_46-68_, *O. vulgaris* RPS15_50-72_, *C. teleta* RPS15_49-71_, *P. pacificus* RPS15_49-71_, *S. pistillata* RPS15_44-66_, *P. syringae* RPS19_50-72_, *Nitrospirae* RPS19_33-55_ and *Aquificae* RPS19_33-55_, and examined their antibacterial activity. As shown in Table 2, all the peptides synthesized exhibited antimicrobial activities against *A. hydrophila* and *S. aureus,* that were basically comparable to or slightly lower than that of BjRPS15_45-67_. All these data suggested that the emergence of the antibacterial activity of RPS15 could be traced to its prokaryotic homolog RPS19.

**Fig 4.**
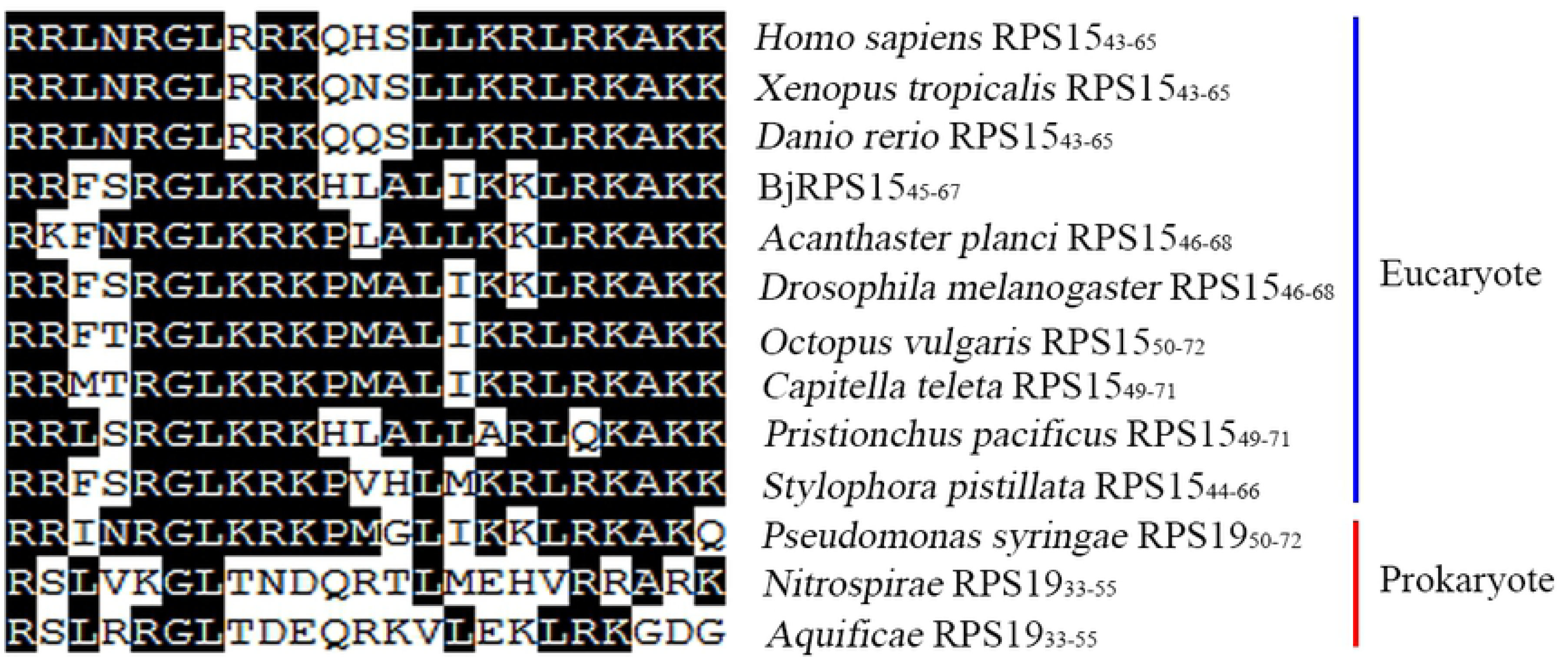
Multiple sequence alignment of the deduced amino acid sequences of BjRPS15_45-67_ counterparts including BjRPS15_45-67_. The identical residues among all the genes are in black.

**Table 2.**
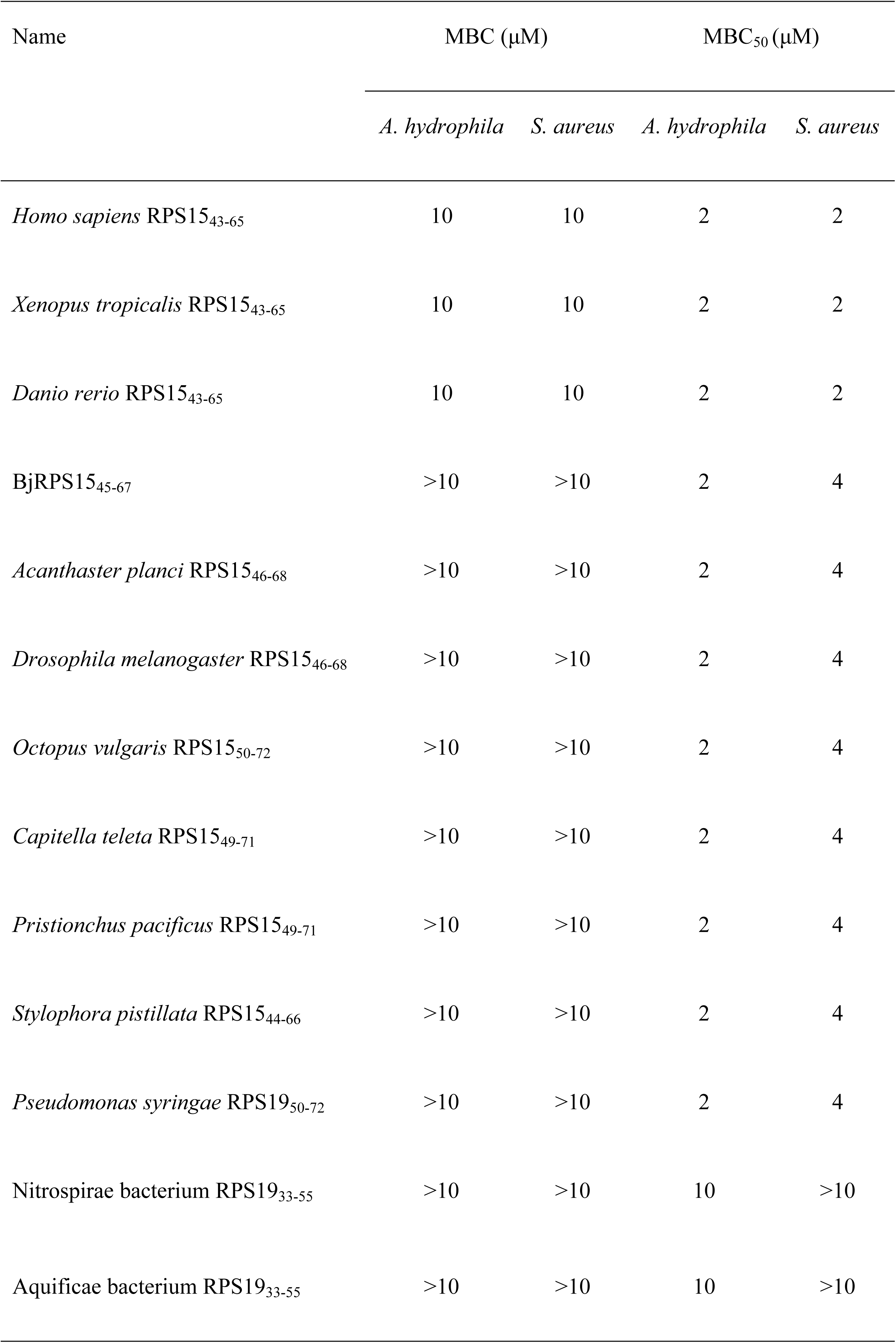
Minimum bactericidal concentration (MBC) and 50% minimum bactericidal concentration (MBC_50_) of BjRPS15 related peptides against the bacteria.

### Destruction of bacterial cells by rBjRPS15 and BjRPS15_45-67_

To examine the effects of rBjRPS15 and BjRPS15_45-67_ on the morphology and structure of bacterial cells, both *A. hydrophila* and *S. aureus* were incubated with rBjRPS15 and BjRPS15_45-67_, and subjected to transmission electron microscopy examination. It was found that both rBjRPS15 and BjRPS15_45-67_ caused a direct damage to the cells of *A. hydrophila* and *S. aureus*, resulting in membrane disruption and cytoplasmic leakage (Fig 5A and B). These indicated that rBjRPS15 and BjRPS15_45-67_ were both bactericidal agents capable of directly killing the bacteria like *A. hydrophila* and *S. aureus*.

**Fig 5.**
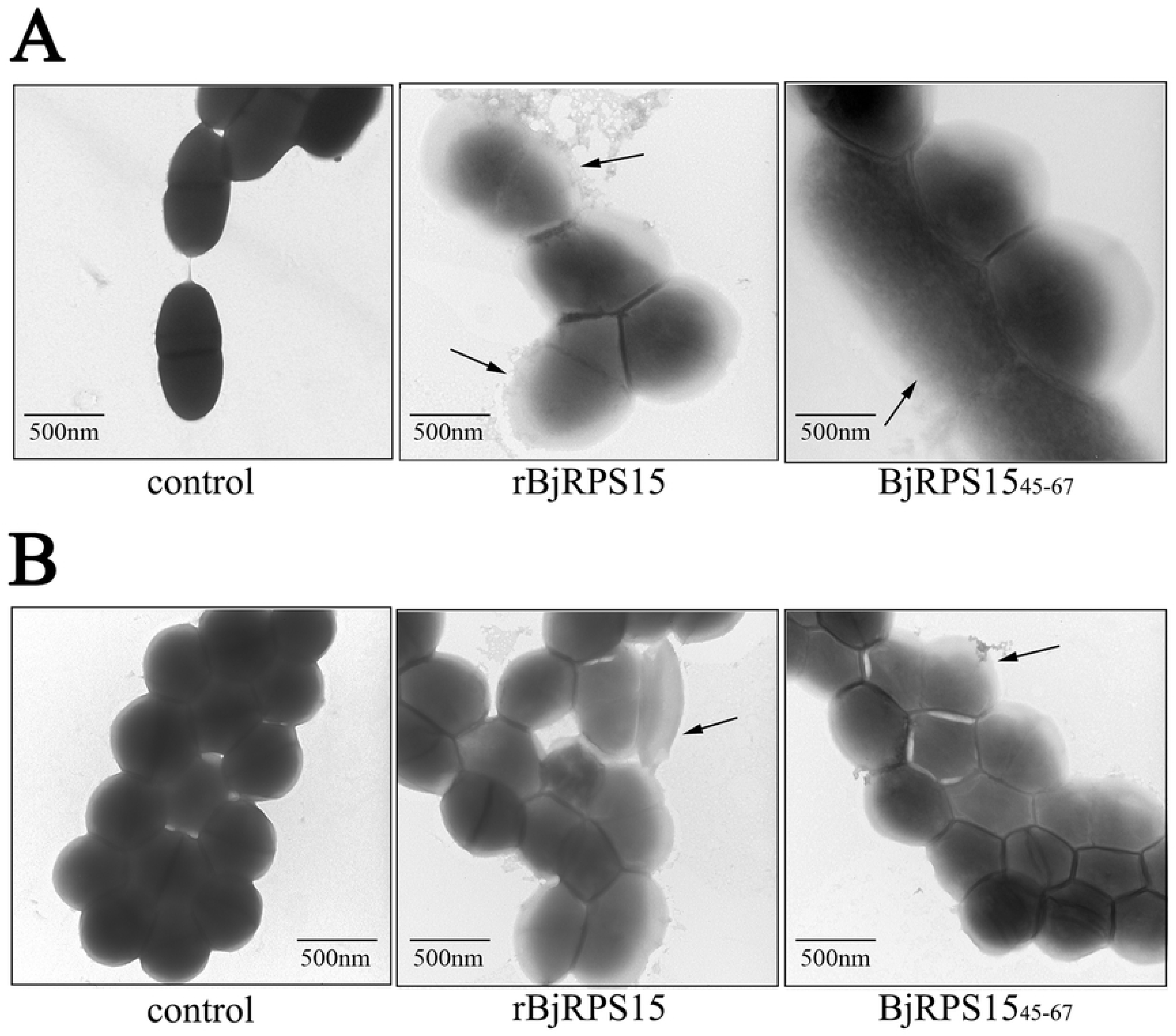
TEM micrographs showing *A. hydrophila* (A) and *S. aureus* (B) that had been exposed to rBjRPS15 (2 μM), BjRPS15_45-67_ (10 μM) or PBS (control) at 25°C for 1 h. The arrows indicated the membrane-disruptive regions of bacteria.

### Bacterial and ligand-binding activities

We then tested if rBjRPS15 could interact with the bacteria. As revealed by Western blotting, rBjRPS15 had strong affinity to *A. hydrophila*, *E. coli*, *S. aureus* and *B. subtilis* (Fig 6A). By contrast, rTRX showed little affinity to the bacteria tested (Fig 6A). These indicated that rBjRPS15 could specifically interact with the Gram-negative and -positive bacteria.

**Fig 6.**
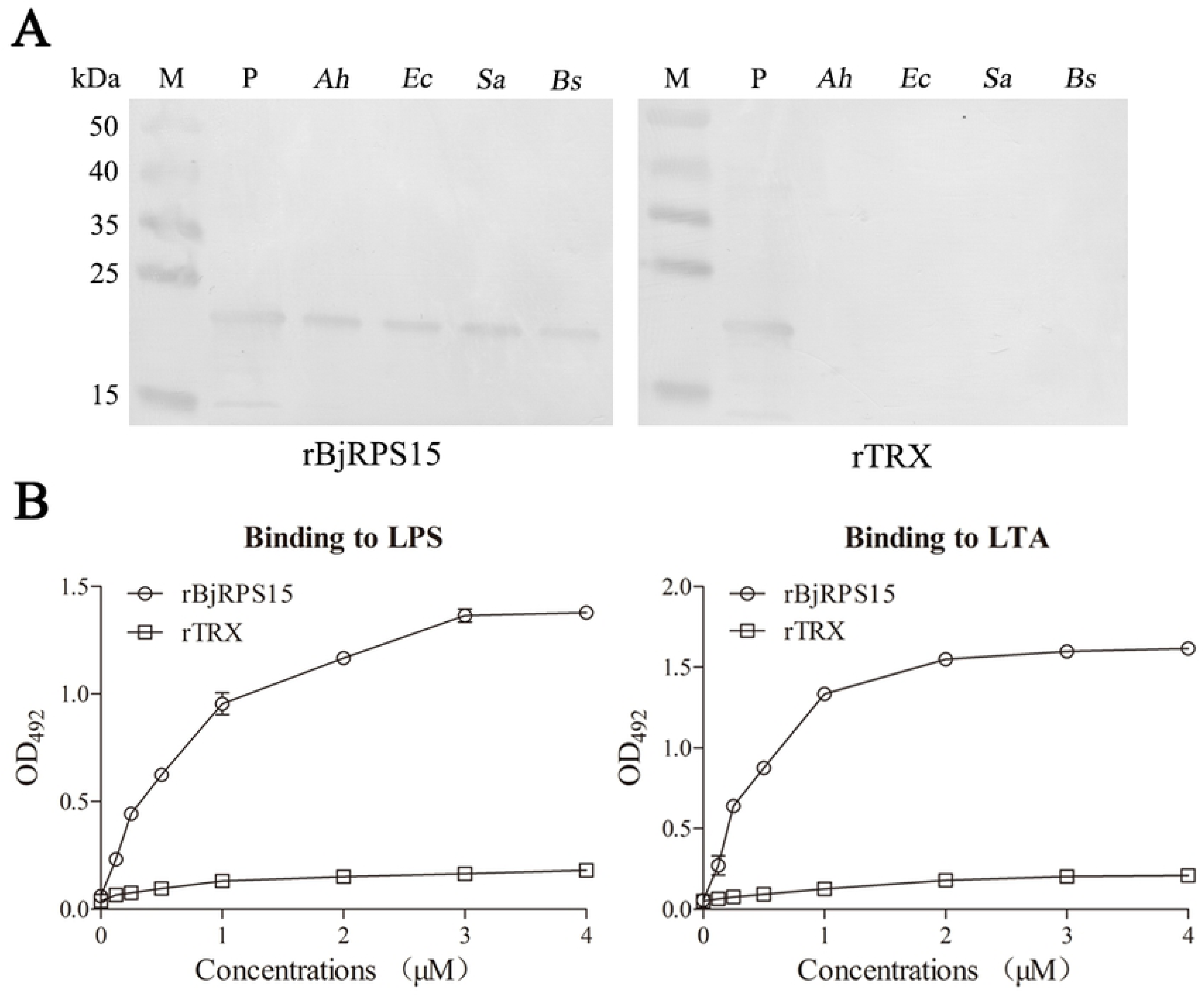
Bacterial binding activity of rBjRPS15 was revealed by Western blotting and ELISA analysis of the affinity of rBjRPS15 to the ligands LPS and LTA. (A) Binding of rBjRPS15 to the bacterial cells. Aliquots of 300 μl of bacterial suspensions (1 × 10^8^ cells/ml) were mixed with 150 μl of 1 μM rBjRPS15 or rTRX (control), respectively. The mixtures were incubated at 25°C for 1 h and washed three times with PBS and re-suspended in 300 μl PBS. The 20 μl bacterial suspensions were subjected to 12% SDS-PAGE gel. The bacterial suspensions on the gels were electroblotted onto PVDF membrane. After incubation with mouse anti-His-tag antibody, the membranes were incubated with the HRP-labeled goat anti-mouse IgG Ab and the bands were visualized using DAB kit. M, molecular mass standards; P, purified recombinant proteins; *Ah*, *Ec*, *Sa* and *Bs* represent *A. hydrophila*, *E. coli*, *S. aureus and A. subtilis* incubated with recombinant proteins, respectively. rTRX was employed as control. (B) ligand binding activity of rBjRPS15. The wells of a 96-well microplate were each coated with one of the ligands and incubated with varying concentrations of the recombinant proteins. After incubation with mouse anti-His-tag antibody, the binding was detected with HRP-labeled goat anti-mouse IgG Ab at 492 nm. Data are shown as mean ± SEM. rTRX was employed as control.

The binding activity of rBjRPS15 to the ligands LPS and LTA was also detected. The results showed that rBjRPS15 was able to bind to LPS and LTA in a dose-dependent manner, whereas rTRX did not (Fig 6B). These indicated that rBjRPS15 interacted with the bacteria via LPS and LTA, suggesting that BjRPS15 might act as multivalent pattern recognition receptors.

### Membrane depolarization by rBjRPS15and BjRPS15_45-67_

The membrane depolarization activities of rBjRPS15 and BjRPS15_45-67_ were assayed using DiSC_3_-5, a potential-dependent distributional fluorescent dye. As shown in Fig 7, the fluorescence intensity of *A. hydrophila*, *E. coli*, *S. aureus* and *B. subtilis* cells treated with rBjRPS15 or BjRPS15_45-67_ was all significantly increased, compared with control (treated with rTRX or PBS). This indicated that rBjRPS15 and BjRPS15_45-67_ both caused depolarization of the bacterial plasma membrane.

**Fig 7.**
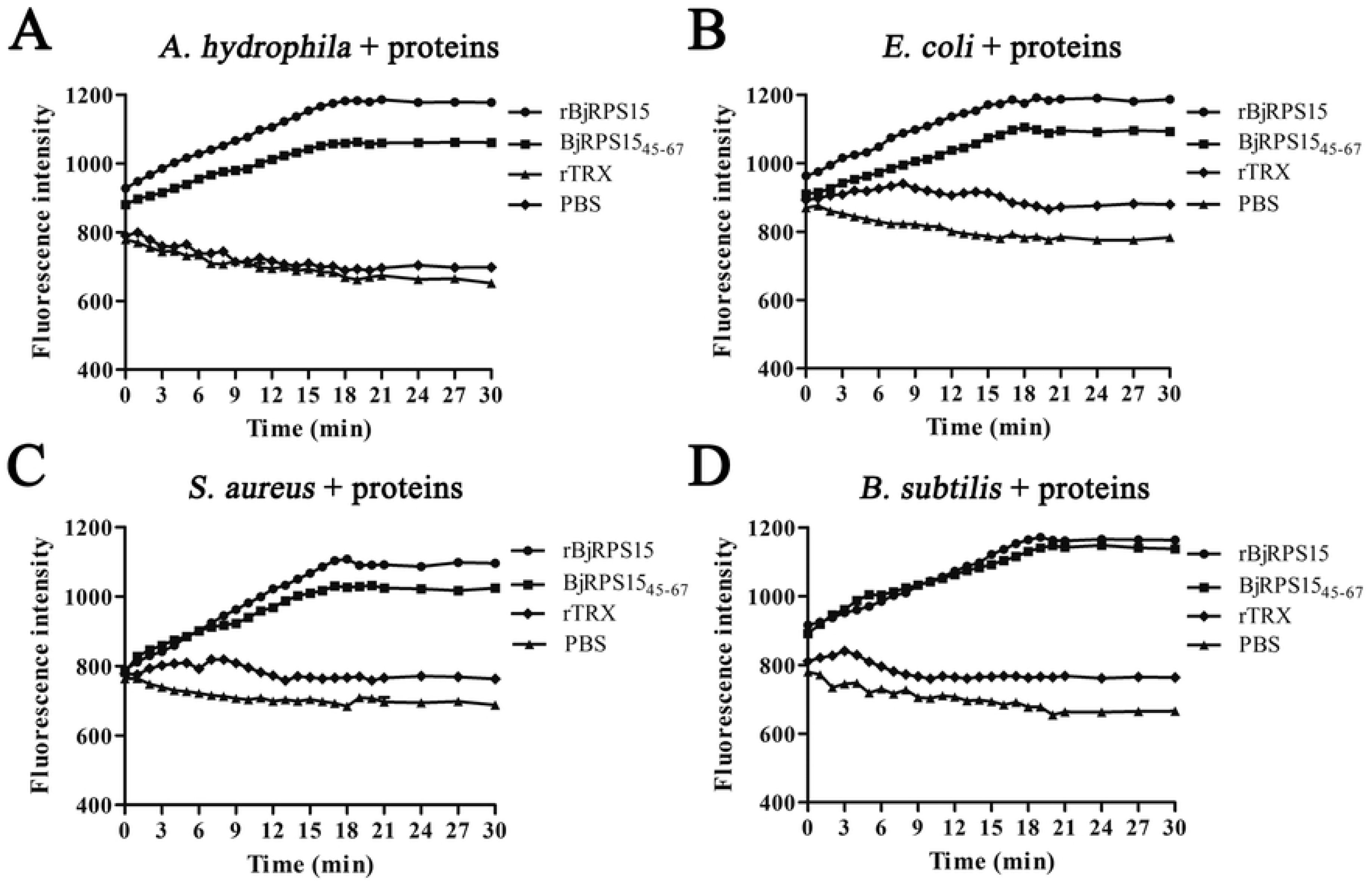
Bacterial membrane depolarization. Depolarization of bacteria cell membranes were detected using DiSC_3_-5 (excitation, 622 nm; emission, 670 nm). rTRX and PBS were used as control.

### Induction of intracellular ROS by rBjRPS15 and BjRPS15_45-67_

High intracellular levels of ROS can cause apoptosis or necrosis. When *A. hydrophila*, *E. coli*, *S. aureus* or *B. subtilis* cells were treated with rBjRPS15 or BjRPS15_45-67_, their intracellular ROS levels were significantly increased (Fig 8). These suggested that both rBjRPS15 and BjRPS15_45-67_ might induce apoptosis/necrosis of the bacterial cells via increased production of intracellular ROS.

**Fig 8.**
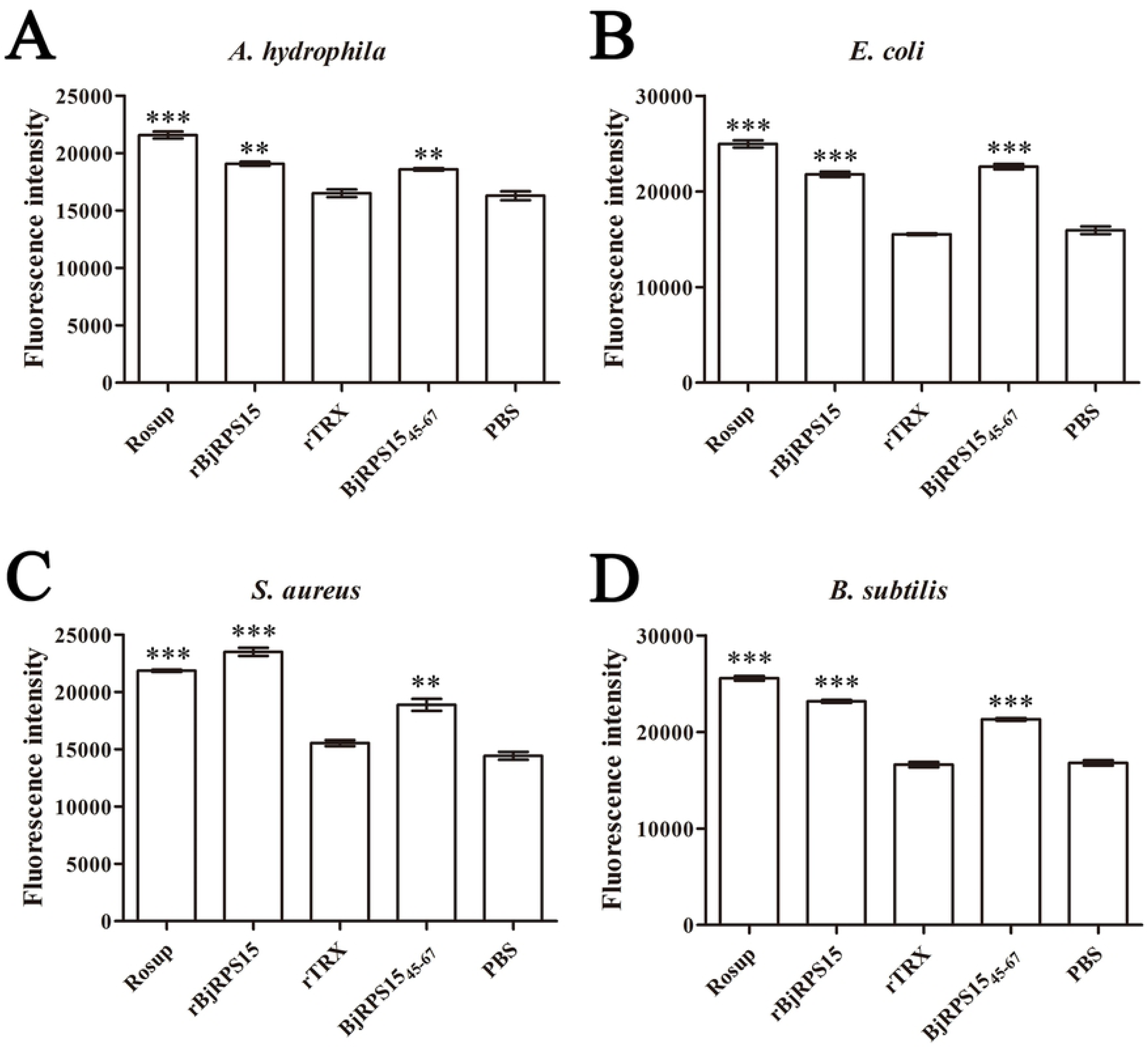
Effects of rBjRPS15 and BjRPS15_45-67_ on intracellular ROS levels. The bacteria *A. hydrophila* (A), *E. coli* (B), *S. aureus* (C) and *B. subtilis* (D) treated with rBjRPS15 and BjRPS15_45-67_. Rosup, a compound mixture, is able to significantly increase ROS levels in cells within 30 min. rTRX and PBS were used as control.

### Non-toxicity of rBjRPS15 and BjRPS15_45-67_ to mammalian cells

To test if rBjRPS15 and BjRPS15_45-67_ were cytotoxic, their hemolytic activities towards human red blood cells (RBCs) were determined. As shown in Fig 9, neither rBjRPS15 nor BjRPS15_45-67_ showed hemolytic activity towards human erythrocytes at all the concentrations tested. By contrast, RBCs incubated with 0.1% Triton X-100, which is usually used as full lysis control, exhibited remarkable hemolysis. The cytotoxicity of rBjRPS15 and BjRPS15_45-67_ to murine RAW264.7 cells was also examined by measuring the cell viability via MTT method. As shown in Table 3, rBjRPS15 and BjRPS15_45-67_ were neither toxic to murine RAW264.7 cells at the concentrations tested. These showed that neither rBjRPS15 nor BjRPS15_45-67_ were toxic to mammalian cells, suggesting that they both showed a high bacterial membrane selectivity.

**Fig 9.**
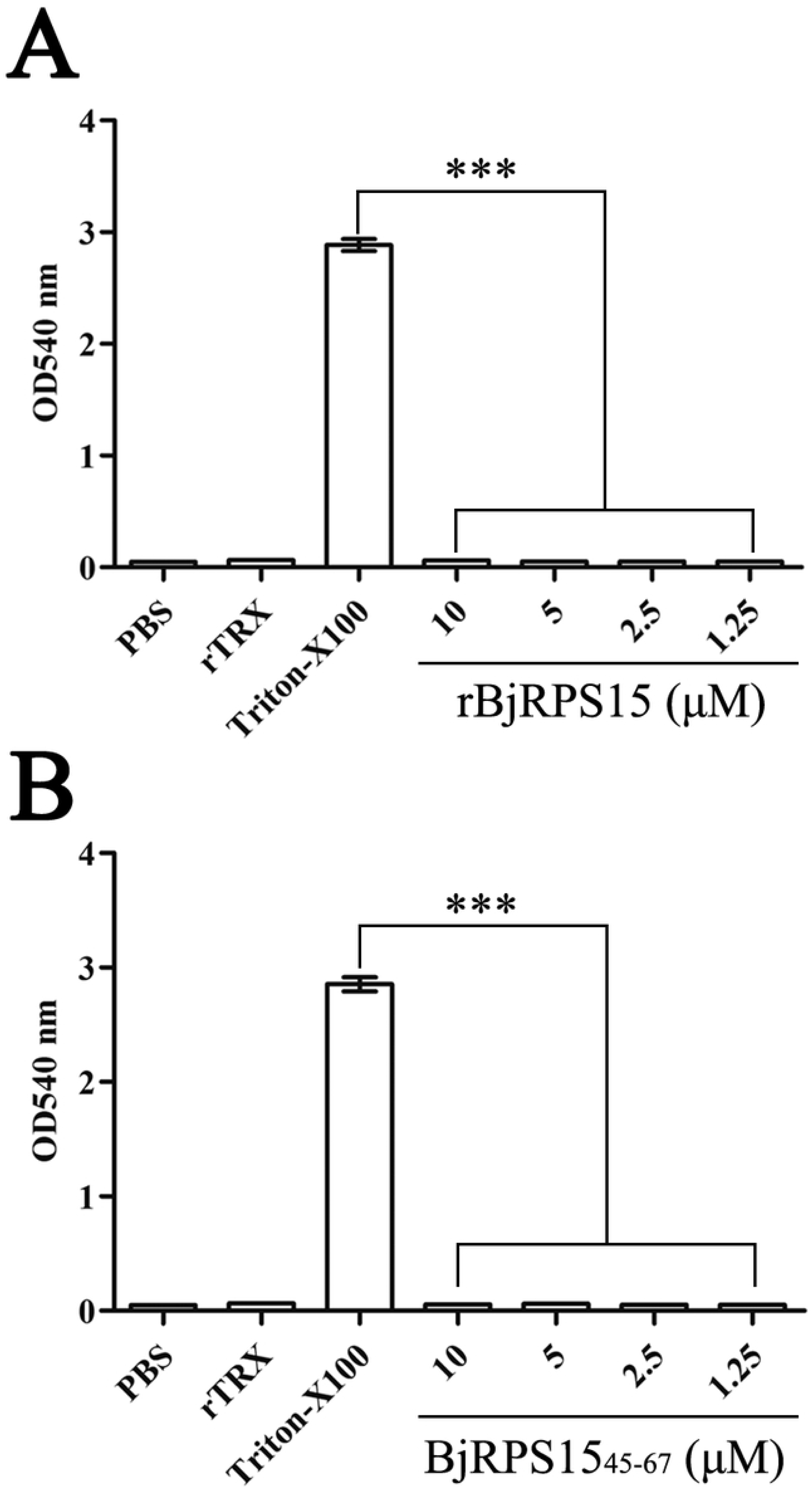
Hemolytic activity of rBjRPS15 and BjRPS15_45-67_ to human blood cells (RBCs). Data were expressed as mean ± SEM (n = 3). The bars represent the standard error of the mean values. The symbol (***) indicates *p* < 0.001 compared with the Triton X-100 treated group. The RBCs incubated with PBS, BSA solution (100 mg/ml), and 0.1% Triton X-100 solution served as blank, negative and positive controls, respectively.

**Table 3.** The percent viability of RAW264.7 cells in the presence of rBjRPS15 and BjRPS15_45-67_. Data were expressed as mean ± SEM (n = 3).

| rBjRPS15 |  | BjRPS15<sub>45-67</sub> |
| --- | --- | --- |

| Concentration (μM) | Viability (%) | Concentration (μM) | Viability (%) |
| --- | --- | --- | --- |
| 0 | 100 | 0 | 100 |
| 1.25 | 102 ± 2 | 1.25 | 106 ± 4 |
| 2.5 | 103 ± 3 | 2.5 | 107 ± 3 |
| 5 | 102 ± 4 | 5 | 110 ± 3 |
| 10 | 105 ± 3 | 10 | 106 ± 5 |

### Presence of extracellular RPS15 in vivo

Next, we examined if extracellular RPS15 was present in animals. Western blotting revealed that the amphioxus humoral fluid was reactive with anti-RPS15 monoclonal antibody, yielding a single band of ∼17 kDa (Fig 10A), well matching the molecular mass predicted by BjRPS15 gene, suggesting the presence of extracellular RPS15 in amphioxus. This was clearly supported by LC/MS/MS analysis (Fig 10B). Similarly, RPS15 was also found to be present in the shrimp hemolymph as well as in zebrafish and mouse sera (Fig 10A). All these indicated that RPS15 existed as extracellular form across widely different animals ranging from invertebrate species to mammals.

**Fig 10.**
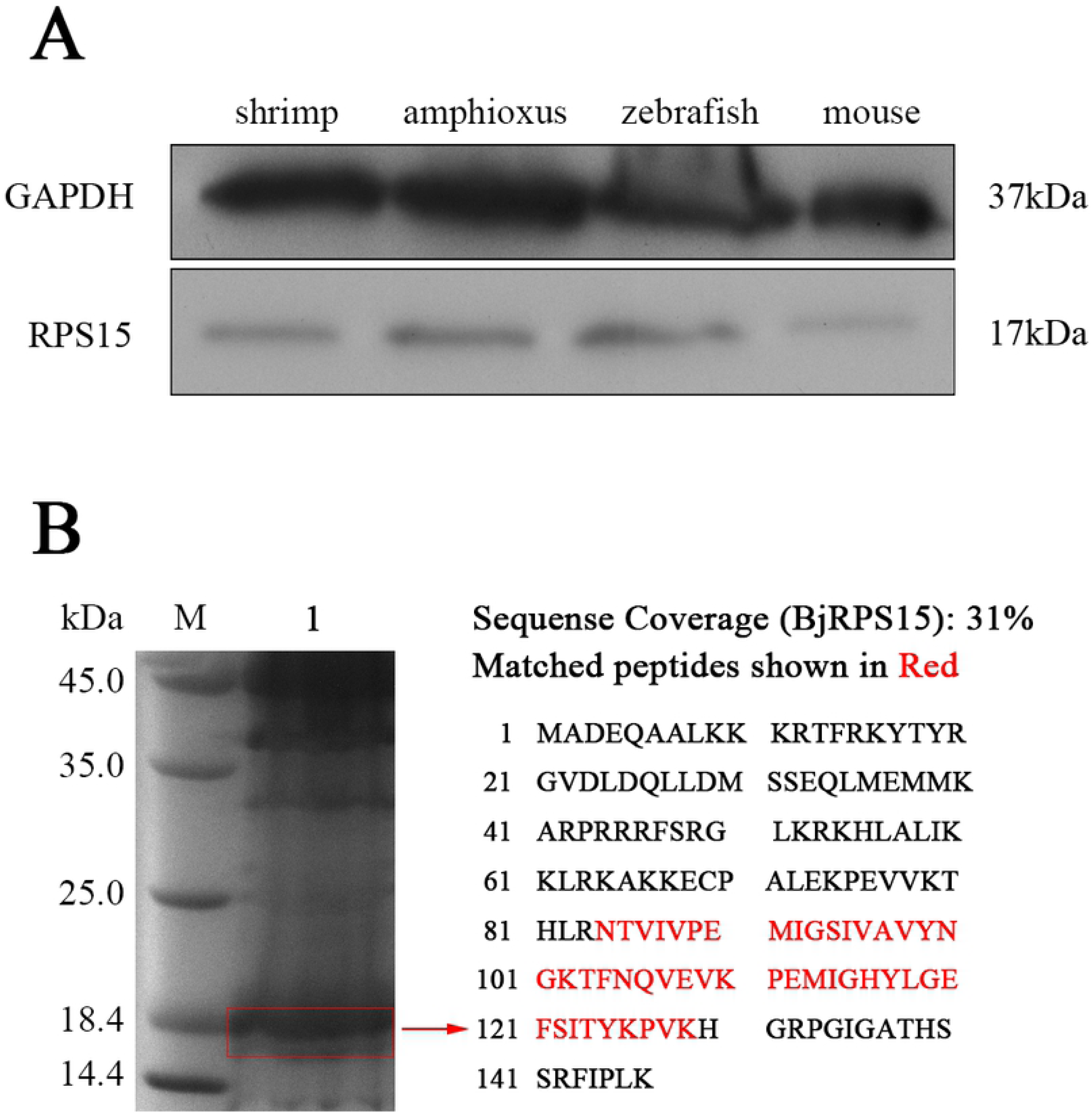
Western blotting and LC/MS/MS analysis. (A) Western blotting analysis of RPS15 in shrimp hemolymph, amphioxus humoral fluids, zebrafish and mouse serums. The shrimp hemolymph, amphioxus humoral fluid, zebrafish and mouse serums were run on a 12% SDS-PAGE gel (Each lane of the gel was loaded with 40 μg of protein). The proteins on the gel were transferred to PVDF membrane. After incubation with anti-RPS15 monoclonal antibody and anti-GAPDH antibody, the membranes were incubated with a secondary HRP-labeled antibody and the bands were visualized by ECL Western blotting substrate. (B) The 40 μg amphioxus humoral fluid loaded on the 12% SDS-PAGE and LC/MS/MS analysis was used to test if RPS15 exist in amphioxus humoral fluid. Lane M, marker; lane 1, amphioxus humoral fluid.

## Discussion

Proteins that fulfil two or more distinct and physiologically relevant biochemical or biophysical functions independent of gene fusions or multiple RNA splice variants are called moonlighting proteins [34, 35]. It is well-known that ribosomal protein S15, or RPS15, a component of the 40S subunit, functions in protein synthesis [36]. In this study, we show for the first time that amphioxus RPS15, BjRPS15, is a previously uncharacterized AMP. It not only functions as a multiple pattern recognition receptor, capable of recognizing LPS and LTA, but also as a bactericidal effector, capable of killing a wide spectrum of bacteria. Potential modes of action of AMPs include interacting with or inserting into bacterial membrane, which can cause lethal depolarization of the usually polarized membrane, scrambling of the normal distribution of lipids between the leaflets of the bilayer, formation of physical pores and loss of critical intracellular targets. We clearly demonstrate that BjRPS15 executes function by a combined action of membranolytic mechanisms including interaction with bacterial membrane through LPS and LTA as well as membrane depolarization. BjRPS15 can also stimulate production of intracellular ROS in bacteria, which may lead to apoptosis/necrosis of the bacterial cells. These indicate that BjRPS15 is a novel member of moonlighting protein, which, in addition to participation in protein synthesis, is involved in anti-infectious response in amphioxus. Intriguingly, RPS15 is present in the humoral fluid of amphioxus, hemolymph of shrimp and sera of zebrafish and mice, indicating that it exists across widely separated taxa. This also has a physiological implication that RPS15 may extracellularly play a critical role in systematic immunity in different animals, protecting them against bacterial infection by interacting with and destructing potential pathogens.

Looking at the amino acid sequences, BjRPS15 shares significantly high identity to its eukaryotic as well as prokaryotic homologs. The 3D structures of BjRPS15 homologs are very similar among both eukaryotes and prokaryotes. Furthermore, the antibacterial features such as high hydrophobic ratio and net positive charge clearly exist in RPS19, an early RPS15 orthologue of bacteria, including *Nitrospirae* sp., *Aquificae* sp. and *P. syringae*. Importantly, it was experimentally proven that the synthesized the peptides of BjRPS15_45-67_ counterparts, including *H. sapiens* RPS15_43-65_, *X. tropicalis* RPS15_43-65_, *D. rerio* RPS15_43-65_, *A. planci* RPS15_46-68_, *D. melanogaster* RPS15_46-68_, *O. vulgaris* RPS15_50-72_, *C. teleta* RPS15_49-71_, *P. pacificus* RPS15_49-71_, *S. pistillata* RPS15_44-66_, *P. syringae* RPS19_50-72_, *Nitrospirae* RPS19_33-55_ and *Aquificae* RPS19_33-55_, all show conspicuous antibacterial activities against the Gram-negative and -positive bacteria tested. These suggest that the antibacterial properties of this family of molecules are very ancient and highly conserved.

An important point about the exploitation of membranolytic antimicrobial therapeutics is that they cannot be cytotoxic to mammalian cell membrane. We find that both BjRPS15 and its truncated BjRPS15_45-67_ are virtually not toxic towards mammalian cells including human erythrocytes and murine macrophages RAW264.7. This denotes that they possess high membrane selectivity to bacterial cells but not to mammalian cells, rendering them ideal lead molecules for the exploitation of new peptide antibiotics against bacteria.

In summary, this present study highlights that RPS15 is a new AMP functioning as a multiple sensor, capable of recognizing LPS and LTA, and an effector, capable of killing the potential pathogens. It also suggests that the antibacterial activities of this family of molecules have ancient origin and high conservation.

## Materials and methods

### Animal culture

All animal experiments performed here conformed to the ethical guidelines established by the Institutional Animal Care and Use Committee of the Ocean University of China. Adult amphioxus *(Branchiostoma japonicum)* collected during the breeding season in the vicinity of Qingdao, China was cultured in aerated seawater at room temperature, and fed twice a day with single-celled algae. Shrimps (*Fenneropenaeus chinensis*) with body weight of approximately 10 to 20 g were purchased from Xiaogang market in Qingdao, and cultured in aerated seawater at room temperature. Wild-type zebrafish (*Danio rerio*) aged 2-3 months purchased from a local fish dealer were maintained at 27 ± 1°C under a controlled light cycle (14 h light/10 h dark) in a 20 L tank with well-aerated tap water, and fed on live bloodworms and Miero Fish Food twice a day. Mice (*Mus musculus*) aged 11 months (no. 20140007) were from Jinan Pengyue Laboratory Animal Breeding Co., Ltd, and housed one per cage in an environmentally controlled atmosphere (temperature 22°C and relative humidity 56%) with a 12 h light/dark cycle. They were given free access to water and diet and provided with shredded wood floor bedding. All the animals were acclimatized for one week before the experiments.

### RNA extraction and cDNAs synthesis

Total RNAs were extracted with Trizol (TaKaRa, China) from *B. japonicum* according to the manufacturer’s instructions. After digestion with the recombinant RNase-free DNase (TaKaRa) to eliminate the genomic contamination, cDNAs were synthesized with reverse transcription kit (TaKaRa) with oligo d(T) primer. The reaction was carried out at 42°C for 50 min and inactivated at 75°C for 15 min. The cDNAs synthesized were stored at −20°C until use.

### Cloning and sequencing of *BjRPS15*

Based on the sequence of *B. belcheri* RPS15 gene (accession number: XP_019635827) in the database of NCBI (http://www.ncbi.nlm.nih.gov/), a pair of primers P1 and P2 (S1 Table) was designed using Primer Premier 5.0 program. The PCR amplification reaction was carried out at 94°C for 5 min, followed by 32 cycles of 94°C for 30 s, 57°C for 30 s and 72°C for 30 s, and a final extension at 72°C for 7 min. The amplification products were gel-purified using DNA gel extraction kit (AXYGEN), cloned into the pGEM-T vector (Invitrogen), and transformed into Trans5ɑ *Escherichia coli* (TransGen). The positive clones were selected and sequenced to verify for authenticity.

### Sequence analysis

The domains and signal peptide of the deduced protein were analyzed by the SMART program (http://smart.embl-heidelberg.de/) and SignalP 5.0 Server (http://www.cbs.dtu.dk/services/SignalP/), respectively. The molecular weight (MW) and isoelectronic points (pI) of the protein were determined by the ProtParam (http://www.expasy.ch/tools/protparam.html). Homology searches in the GenBank database were carried out by BLAST server (http://www.ncbi.nlm.nih.gov/BLAST/). The information of exon-intron organization was obtained from NCBI database. Multiple protein sequences were aligned using the MegAlign program of the LASERGENE software suite (DNASTAR). The SWISS-MODEL prediction algorithm (https://swissmodel.expasy.org/) was applied to generate the three-dimensional (3D) structure model. CAMP server (http://www.camp.bicnirrh.res.in/predict/) was used to predict the core sites for antimicrobial activity, and Antimicrobial Peptide Calculator and Predictor at APD (http://aps.unmc.edu/AP/main.php) used to calculate the total hydrophobic ratio and net charge.

### Quantitative real-time PCR (qRT-PCR)

qRT-PCR was used to examine the transcriptional profile of *BjRPS15* in the different tissues of *B. japonicum*, including the hepatic caecum, hind-gut, gill, muscle, notochord, testis and ovary, as described by Wang et al. [14] and Yang et al. [15]. The PCR primer pairs P3 and P4 as well as P5 and P6 (S1 Table) specific of *BjRPS15* and *EF1α* were designed using primer 5.0 program. The *EF1α* gene was chosen as the reference for internal standardization. The expression level of *BjRPS15* relative to that of *EF1α* gene was calculated by the comparative Ct method (2^−ΔΔCt^) [16].

The qRT-PCR was also performed to assay the transcriptional profile of *BjRPS15* in response to challenge with the bacteria *Aeromonas hydrophila* (ATCC 35654), *E. coli* (ATCC 25922), *Staphylococcus aureus* (ATCC 25923) and *Bacillus subtilis* (ATCC 6633) as well as the bacterial signature molecules lipopolysaccharide (LPS) and lipoteichoic acid (LTA) as described by Wang et al. [14]. The *B. japonicum* were cultured in 1 L of sterilized seawater containing 10^8^ cells/ml of *A. hydrophila*, *E. coli*, *S. aureus* or *B. subtilis* or 10 μg/ml of the bacterial signature molecules LPS (Sigma, USA) or LTA (Sigma, USA) [14,17,18], and sampled at 0, 2, 4, 8, 12, 24, 48, and 72 h after the exposure. Extraction of total RNAs, cDNA synthesis and qRT-PCR were carried out as above.

### Construction of expression vector

The sequence encoding BjRPS15 was amplified by PCR using the primer pairs P7 and P8 (S1 Table) with *EcoR* I and *Xho* I sites in the forward and reverse primers, respectively. The PCR products were sub-cloned into the plasmid expression vector pET-28a (Novagen) previously cut with the restriction enzymes *EcoR* I and *Xho* I. The identity of inserts was verified by sequencing, and the constructed plasmid was designated *pET-28a/BjRPS15*.

### Expression and purification of rBjRPS15

The plasmid *pET28a/BjRPS15* was transformed into *E. coli* transetta (DE3) cells. Induced expression and purification of recombinant BjRPS15, rBjRPS15, were performed as described by Gao et al. [19]. Recombinant thioredoxin His Tag (rTRX) used for control was similarly prepared. The purity of the eluted samples and purified proteins were analyzed by a 12% SDS-PAGE gel and stained with Coomassie brilliant blue R-250. The concentrations of the recombinant proteins were determined by BCA method.

### Western blotting

Western blotting was conducted as described by Yao et al. [20]. The anti-His-tag mouse monoclonal antibody (CWBIO) used was diluted 1:4000 with 4% BSA in PBS.

### Antimicrobial activity assay

The antimicrobial activity of rBjRPS15 was assayed by the method of Shi et al. [21]. Briefly, aliquots of 50 μl of *A. hydrophila*, *E. coli*, *S. aureus* or *B. subtilis* suspension (3 × 10^4^ cells/ml) were each mixed with 50 μl of rBjRPS15 solution (0, 0.5, 1, 2, 3, and 4 μM) and incubated at 25°C for 1 h. Each of the bacterial mixtures was then plated onto 3 agar plates (30 μl/plate). After incubation at 37°C for 12 h, the resulting bacterial colonies in each plate were counted. The percent of bactericidal activity was calculated as follows: [number of colonies (control-test)/number of colonies (control)] × 100 (n = 3). The rTRX was used as control.

The core site of BjRPS15 for antimicrobial activity was predicted by CAMP server, and the residues positioned at 45-67 (RRFSRGLKRKHLALIKKLRKAKK), designated BjRPS15_45-67_, were identified as the only candidate. Therefore, BjRPS15_45-67_ was synthesized by Shanghai Sangon Biological Engineering Technology & Services Co., Ltd, using standard solid-phase FMOC method, and used for antimicrobial activity assay. Because C-terminal amidation is common in antimicrobial peptides (AMPs) [22], the C-termini of the peptide was thus amidated. The peptide synthesized was purified to >95% by high-performance liquid chromatography (HPLC) and the mass of the synthetic peptide was verified by mass spectrometer (lcms-2010a, Shimadzu, Japan). The peptide was dissolved in PBS (2 mg/ml) and stored at −80°C till used. The antimicrobial activity of BjRPS15_45-67_ (0, 0.625, 1.25, 2.5, 5, 10 μM) was determined as described above.

Sequence alignment revealed that BjRPS15 shared more than 59.7% identity to eukaryotic RPS15 and more than 41.2% identity to its prokaryotic homolog RPS19 (S1 Fig). Thus, we synthesized the core regions (corresponding to the region BjRPS15_45-67_) of RPS15 of the eukaryotes including *Homo sapiens, Xenopus tropicalis, Danio rerio*, *Acanthaster planci, Drosophila melanogaster, Octopus vulgaris, Capitella teleta, Pristionchus pacificus* and *Stylophora pistillata* as well as those of RPS19 of the prokaryotes including *Pseudomonas syringae, Nitrospirae* sp. and *Aquificae* sp. (see Fig 4 below) to test the conservation of RPS15 antibacterial activity. The antibacterial activities of the synthesized peptides were examined as above.

### Transmission electron microscopy (TEM)

TEM was performed to test the effect of rBjRPS15 and BjRPS15_45-67_ on the morphology and structures of the bacteria *A. hydrophila* and *S. aureus* as described by Liu et al. [23]. Briefly, aliquots of 500 μl of the bacterial suspensions containing 5 × 10^7^ cells/ml were mixed with 500 μl of rBjRPS15 (4 μM) or BjRPS15_45-67_ (20 μM), respectively. In parallel, aliquots of 500 μl of the bacterial suspensions were mixed with 500 μl PBS as control. The mixtures were incubated at 25°C for 1 h, fixed in 2.5% glutaraldehyde in 100 mM PBS, and then dropped onto 400-mesh carbon-coated grids and allowed to stand at room temperature for 3 min for negative staining. Excess fluid was removed by touching the edge of filter paper. The grids were then put into 2% phosphotungstic acid for 3 min, dried by filter paper, and observed under a JEOL JSM-840 transmission electron microscope.

### Bacterial binding assay

To test the bacterial binding activity of rBjRPS15, the bacteria *A. hydrophila*, *E. coli*, *S. aureus* and *B. subtilis* were cultured to mid-logarithmic phase, and collected by centrifugation at 6000 g for 5 min. After washing twice with PBS, the bacteria were re-suspended in PBS, giving a density of 1 × 10^8^ cells/ml. Aliquots of 300 μl of bacterial suspensions were mixed with 150 μl of 1 μM rBjRPS15 or rTRX (control), respectively. The mixtures were incubated at 25°C for 1 h, and centrifuged at 6000 g at 4°C for 5 min. The bacterial pellets were washed three times with PBS and re-suspended in 300 μl PBS. The bacterial suspensions were subjected to 12% SDS-PAGE gel and the binding activity was determined by Western blotting as described above.

### Ligand binding assay

An enzyme-linked immunosorbent assay (ELISA) was preformed to examine the binding of rBjRPS15 to the ligands LPS and LTA. Aliquots of 50 μl of 40 μg/ml LPS or LTA were applied to each well of a 96-well microplate and air-dried at 25°C overnight. The plates were incubated at 60°C for 30 min to fix the ligands, and then each well was blocked with 100 μl of 1 mg/ml BSA in PBS at 37°C for 2 h. After washing four times with PBST, a total of 100 μl PBS containing 0.1 mg/ml BSA and different concentrations (0, 0.0625, 0.125, 0.25, 0.5, 1, 1.5 and 2 μM) of rBjRPS15 or rTRX (control) was added into each well and incubated at 25°C for 3 h. The wells were rinsed four times with PBST, and incubated with 100 μl of mouse anti-His-tag antibody (CWBIO), diluted 1:5000 with 4% BSA in PBS, at 37°C for 1 h. After washing four times with PBST, the wells was then incubated with 100 μl of HRP-labeled goat anti-mouse IgG Ab (CWBIO), diluted 1:8000 with 4% BSA in PBS, at room temperature for 1 h. Subsequently, the wells were washed four times with PBST, added with 75 μl of 0.4 mg/ml O-phenylenediamine (Amresco) in the buffer consisting of 51.4 mM Na_2_HPO_4_, 24.3 mM citric acid and 0.045% H_2_O_2_ (pH5.0), and reacted at 37°C for 10 min. Finally, 25 μl of 2 M H_2_SO_4_ was added into each well to terminate the reaction, and absorbance at 492 nm was monitored by a microplate reader (GENios Plus; Tecan).

### Membrane depolarization assay

The assay for membrane depolarization activity of rBjRPS15 and BjRPS15_45-67_ was performed with the membrane potential-sensitive dye 3,3’-dipropylthiacarbocyanine iodide (DiSC_3_-5; Sigma) and the bacteria *hydrophila*, *E. coli*, *S. aureus* and *B. subtilis*, according to the method of Lee et al. [24]. The bacterial cells in the mid-logarithmic phase were harvested by centrifugation at 6000 g for 10 min, washed in 5 mM HEPES buffer (pH7.3) containing 20 mM glucose, and re-suspended in 5 mM HEPES buffer containing 20 mM glucose and 100 mM KCl to an OD_600_ of 0.05. Aliquots of 100 μl of the bacterial suspensions, supplemented with 0.5 μM DiSC_3_-5, were applied to each well of a 96-well flat bottom white microplate, and allowed to stand for 30 min at room temperature to get a steady baseline of fluorescence intensity. The bacterial suspensions were then mixed with 100 μl of PBS containing 2 μM rBjRPS15, 2 μM rTRX, 10 μM BjRPS15_45-67_ or PBS alone. rTRX and PBS were used as control. Changes in the fluorescence intensity were continuously recorded for 30 min with a TECAN-GENios plus spectrofluorimeter at an excitation wavelength of 622 nm and an emission wavelength of 670 nm.

### Reactive oxygen species assay

The levels of reactive oxygen species (ROS) were measured as described by Cui et al. [25]. The bacteria *A. hydrophila*, *E. coli*, *S. aureus* and *B. subtilis* were re-suspended in the corresponding culture medium containing 10 μM DCFH2-DA, yielding a density of 10^7^ cells/ml. After incubation at 37°C for 30 min, the bacteria were collected by centrifugation at 6000 g at room temperature for 10 min. The bacterial cells were washed three times with PBS, re-suspended in 1 ml of PBS containing 1 μM rBjRPS15, 1 μM rTRX or 5 μM BjRPS15_45-67_. For positive control, the bacterial cells were re-suspended in 1 ml PBS containing 50 μg/ml Rosup, a compound mixture, that can significantly increase ROS levels in cells within 30 min. For blank control, the cells were re-suspended in 1 ml PBS alone. The bacterial suspensions were incubated at 25°C for 1 h and the fluorescence intensities were recorded immediately with a TECAN-GENios plus spectrofluorimeter at an excitation wavelength of 488 nm and an emission wavelength of 525 nm.

### Hemolytic activity assay

Human red blood cells (RBCs) were used to test the hemolytic activity of rBjRPS15 and BjRPS15_45-67_ as described by Hu et al. [26]. Healthy human blood was obtained and placed in an EDTA anticoagulant tube. The RBCs were collected by centrifugation at room temperature at 1000 g for 10 min. After washing three times with PBS, the RBC pellets were suspended in PBS to give a concentration of 4% (v/v). Aliquots of 200 μl RBCs suspension were mixed with 200 μl of different concentrations of rBjRPS15 (0, 1.25, 2.5, 5 and 10 μM) or BjRPS15_45-67_ (0, 1.25, 2.5, 5 and 10 μM), respectively. After incubation at 37°C for 1 h, the mixtures were centrifuged at room temperature at 1000 g for 10 min. The supernatants were collected and added into a 96-well plate. The absorbance was measured at 540 nm under a microplate reader (Multiskan GO; Thermo Scientific). RBCs incubated with PBS, rTRX solution (10 μM), and 0.1% Triton X-100 solution served as blank, negative and positive controls, respectively.

### 3-(4,5-dimethylthiazol-2-yl)-2,5-diphenyltetrazolium bromide (MTT) assay

To test if rBjRPS15 and BjRPS15_45-67_ are cytotoxic to murine RAW264.7 cells, MTT assay was performed as described by Hu et al. [26]. RAW264.7 cells were suspended in serum-free DMEM and aliquots of 180 μl of the cell suspension (1 × 10^6^ cells/ml) were sampled into a 96-well plate and cultured at 37°C with 5% CO_2_ for 2 h. After removal of the medium, aliquots of 200 μl serum-free DMEM with different concentrations of rBjRPS15 (0, 1.25, 2.5, 5 and 10 μM) and BjRPS15_45-67_ (0, 1.25, 2.5, 5 and 10 μM) were added to each well of a 96-well microplate, incubated for 4 h, and then 20 μl of MTT solution (5 mg/ml) was added into each well. After incubation for another 4 h, the medium was removed and 150 μl of dimethyl sulfoxide (DMSO) was added. The absorbance at 492 nm was measured under a microplate reader. For control, the solution of rBjRPS15 and BjRPS15_45-67_ was substituted by PBS alone, and the assays were similarly processed. The percent viability against the control was calculated as follows: (OD of treated groups/OD of control groups) × 100 (n = 3).

### Assay for extracellular RPS15 in vivo

Western blotting was used to test if RPS15 was present in the humoral fluid of amphioxus, hemolymph of shrimp and sera of zebrafish and mouse. The humoral fluid was prepared by the method of Pang et al. [27], the hemolymph prepared from shrimps by the method of Zhang et al. [28], and the sera of zebrafish and mouse were prepared as described by Babaei et al. [29] and Greenfield [30]. Western blotting was performed as described above using anti-RPS15 monoclonal antibody (1:1000; Abcam, UK) and anti-GAPDH antibody (1:3000; Bioss, China) as the primary antibodies, respectively, and HRP-labeled antibody (1:6000; CWBIO, China) as the secondary antibody. The bands were visualized by ECL Western blotting substrate (Thermo Fisher Scientific, USA).

To further verify the presence of BjRPS15 in the humoral fluid of amphioxus, the liquid chromatography-tandem mass spectrometry (LC/MS/MS) analysis was performed. The humoral fluid was run on a 12% SDS-PAGE gel and stained with Coomassie Brilliant Blue R-250. The band with an expected molecular weight of ∼17 kDa corresponding that of RPS15 was cut off and subjected to LC/MS/MS analysis (Beijing Protein Innovation Co., Ltd, China).

### Statistical analysis

The experiments were performed in triplicate, and repeated three times. Data were subjected to statistical evaluation with unpaired t-test, and the value *p*<0.05 was considered as significant. All the data were expressed as mean ± SEM.

## Acknowledgements

The authors acknowledge the substantive input from all members of the Laboratory for Evolution & Development. This work was supported by the Ministry of Science and Technology (MOST) of China (2018YFD0900505) and the Marine S&T Fund of Shandong province for Pilot National Laboratory for Marine Science and Technology (Qingdao) (2018SDKJ0302-1).

## Supporting information

**S1 Fig.** The sequence identity of BjRPS15. The amino acid identity and divergence was calculated using the Clustal W program within the MegAlign of the DNASTAR software package (version 5.0). Percent identity compares sequences directly, without accounting for phylogenetic relationships. Divergence is calculated by comparing sequence pairs in relation to the phylogeny reconstructed by MegAlign.

**S2 Fig.** The 3D structures of BjRPS15_45-67_ counterparts derived from eukaryotes and prokaryotes.

**S1 Table.** Sequences of the primers used in this study.

**S2 Table.** Accession numbers of RPS15 proteins used in multiple alignment and sequence identity analysis.

